# Integrating sex-bias into studies of archaic admixture on chromosome X

**DOI:** 10.1101/2022.08.30.505789

**Authors:** Elizabeth T. Chevy, Emilia Huerta-Sánchez, Sohini Ramachandran

## Abstract

Evidence of interbreeding between archaic hominins and humans comes from methods that infer the locations of segments of archaic haplotypes, or ‘archaic coverage’ using the genomes of people living today. As more estimates of archaic coverage have emerged, it has become clear that most of this coverage is found on the autosomes— very little is retained on chromosome X. Here, we summarize published estimates of archaic coverage on autosomes and chromosome X from extant human samples. We find on average 7.9 times more archaic coverage on autosomes than chromosome X, and identify broad continental patterns in this ratio: greatest in American samples, and least in South Asian samples. We also perform extensive simulation studies to investigate how the amount of archaic coverage, lengths of coverage, and rates of purging of archaic coverage are affected by sex-bias caused by an unequal sex ratio within the archaic introgressors. Our results generally confirm that, with increasing male sex-bias, less archaic coverage is retained on chromosome X. Ours is the first study to explicitly model such sex-bias and its potential role in creating the dearth of archaic coverage on chromosome X.

**Author summary:** Tens of thousands of years ago, humans interbred with our close hominin relatives (*e*.*g*. Neanderthals), which we know from finding segments of archaic hominin DNA in our genomes. Up to 4% of a human genome may be archaic DNA, but most of that archaic part is on the autosomes (the non-sex chromosomes). Chromosome X usually contains 3 to 10 times less archaic DNA than the autosomes. Also unlike the autosomes, it is always passed down by mothers, but only sometimes by fathers. There are several hypotheses for why chromosome X has less archaic DNA than the autosomes; one that has not been fully explored is whether the archaic hominins that interbred with our ancestors were mostly male or mostly female, known as ‘sex-bias’. In this paper, we use simulations to investigate whether sex-bias could produce less archaic DNA on chromosome X. Using simulation studies, we find that when the archaics are mostly male, modern humans end up with less archaic DNA on chromosome X than their autosomes, compared to when there is a female-bias or no sex-bias. Therefore, male sex-bias could be contributing to the difference in the amount of archaic DNA on chromosome X versus the autosomes. Of course, there are still plenty of other factors to be explored about how demography and selection have shaped our DNA. Studying patterns like this helps us learn more about early hominin natural history, and contextualizes archaic interbreeding events among other sex-biased events in human history.

## Introduction

Secondary contact between archaic hominin (*e*.*g*., Neanderthal) and modern human groups around 50,000 years before present generated individuals with genomes containing a mixture of both archaic and modern human genetic material, and genomic evidence of this archaic genetic material survives in the human gene pool today. Recent computational methods [1–3] enable inferring which ‘tracts’ (or, segments of haplotypes) of an individual’s genome are retained from an archaic introgression event. We refer to the proportion of an individual’s genome, or of a given chromosome of interest, contained within archaic tracts as that individual’s ‘archaic coverage’. Among extant modern humans who harbor some archaic genetic material, archaic coverage across the autosomes typically ranges between 1-3% [3]. However, as Fig 1 demonstrates, the archaic coverage observed on chromosome X is much less than that observed on the autosomes [1–5].

**Fig 1.**
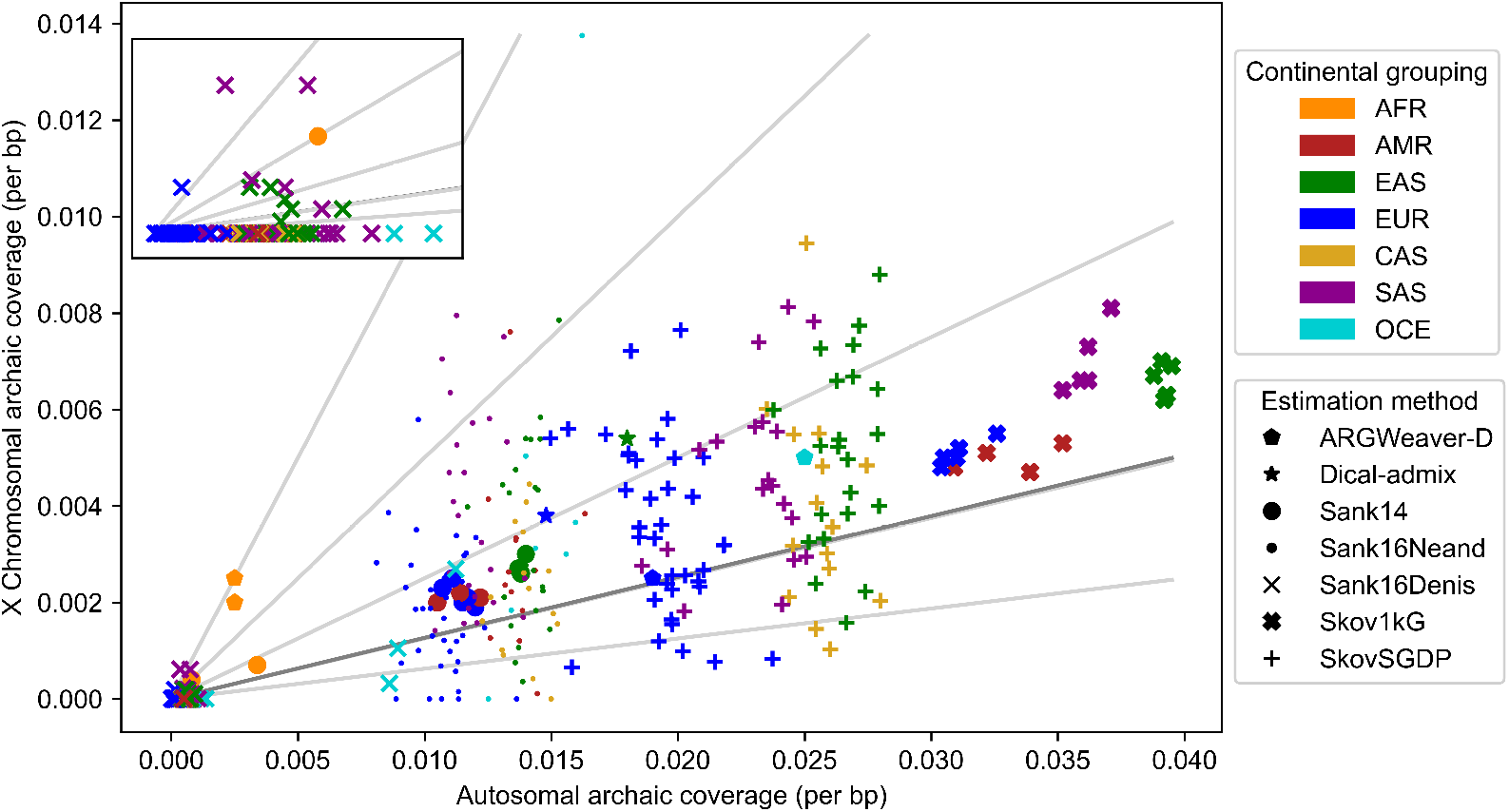
Summary of published empirical estimates of archaic coverage in global groups show greater coverage rates on autosomes than chromosome X. Each point indicates the per-bp amount of archaic coverage on autosomes and chromosome X in a global human group; see S1 Table for numerical values. Colors indicate continental group; see Table 1 for key. Shapes indicate estimation method; see S2 Table for details. Light grey clines show autosomal to chromosome X coverage ratios of 1 (*i*.*e*., equal per bp coverage), 2 (*i*.*e*., autosomal archaic coverage is double that of chromosome X), 4, 8, and 16. Dark grey cline shows median autosomal to chromosome X coverage ratio of 7.91. Nearly all points are well below the 1:1 line, indicating greater per-bp coverage on autosomes than chromosome X.

Despite the scientific attention paid to autosomal archaic coverage in modern humans, the precise factors behind the archaic coverage discrepancy between chromosome X and autosomes remain unresolved. Many studies have examined how different modes of selection and introgression may have reduced archaic coverage to the levels observed in modern humans living today [1, 2, 6–11]. Of these, only a few [1, 2, 6, 8] have attempted to study how evolutionary processes influence archaic coverage on chromosome X. Sankararaman et al. [1] posited that the large-X effect, an excess of hybrid incompatibility loci on chromosome X, reduced male fitness in admixed individuals and consequently drove archaic variants from these X-linked loci. Still fewer studies [2, 6, 8] have modeled the unique inheritance pattern of chromosome X, thereby acknowledging that X-linked mutations can be differentially exposed to selection in males versus females. These studies have argued that background selection against archaic variants is sufficient to explain the depletion of chromosome X archaic coverage.

**Table 1.**
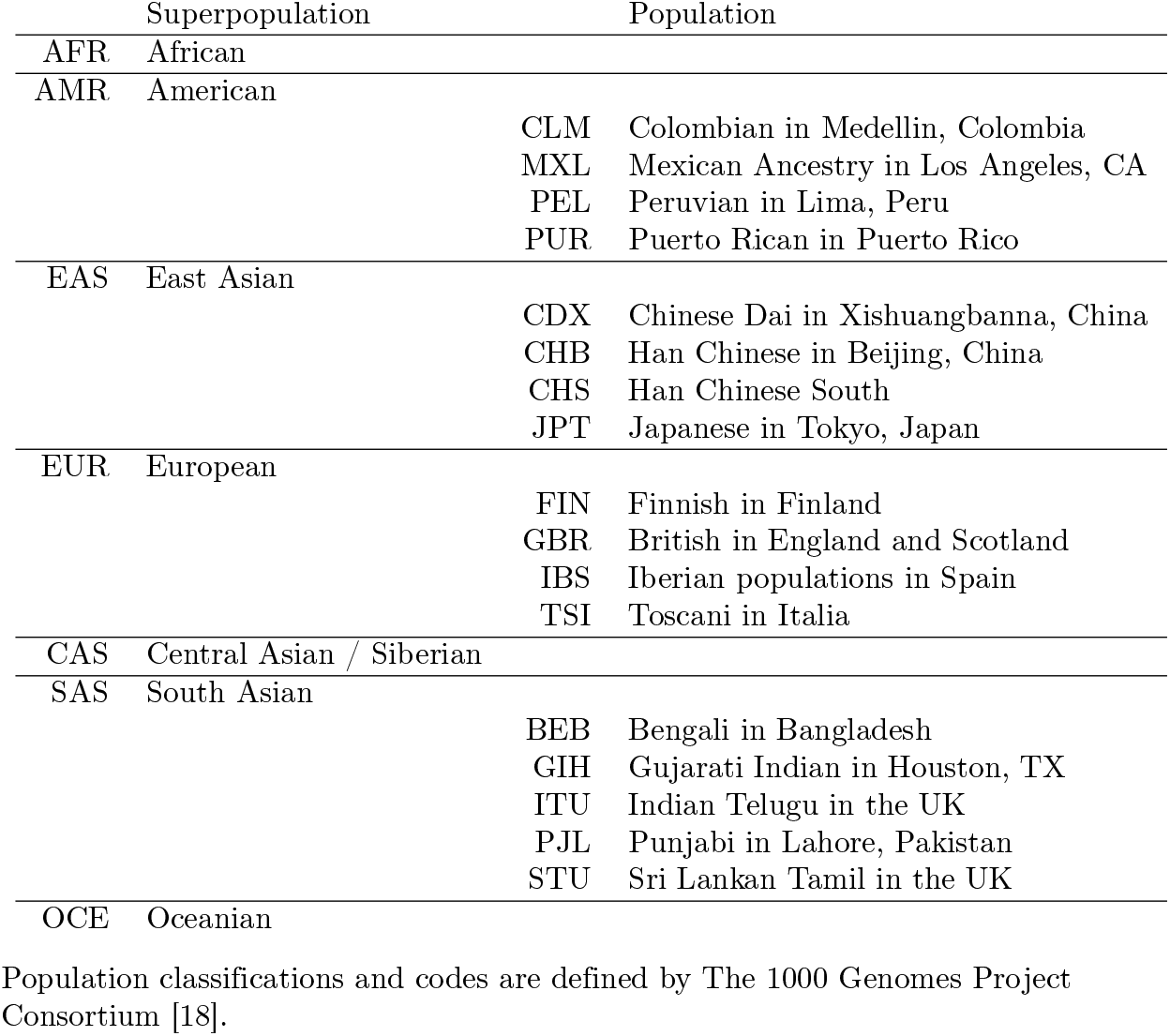
Population abbreviations used in Figs 1 and 2.

Sex-bias—a phenomenon that is readily observed in human societies today [12–14], as well as in historical introgression events [15, 16]—could also drive the observed discrepancy in archaic coverage between chromosome X and the autosomes. However, to our knowledge, no study has yet considered how sex-biased demography (unequal representation of males and females in demographic processes such as migration or population founding events) might interact with sex-specific chromosome X inheritance and natural selection to produce the paucity of archaic coverage observed today on chromosome X. For example, male-bias in the archaic hominin donor pool could result in lower archaic coverage on chromosome X in modern humans today. If male archaics disproportionately contributed deleterious archaic variation, recessive variants would be more efficiently purged from chromosome X than the autosomes, due to the hemizygous state of chromosome X in males. Even for neutral archaic variation, a disproportionate contribution from males would leave less archaic coverage on chromosome X due to the lower copy number of chromosome X in males than females.

In this study, we test whether sex-bias in archaic-human introgression events may have contributed to relatively low archaic coverage on chromosome X versus autosomes in modern humans, and how sex-bias can interact with dominance to produce the observed discrepancy in archaic coverage. To do so, we simulate an archaic introgression scenario using either chromosome X or autosomal inheritance using SLiM3 [17]. We calculate the archaic coverage observed in simulated haplotypes at the present day (end of the simulation), and investigate how dominance and sex-bias affect the amount, distribution, and temporal dynamics of archaic coverage on chromosome X.

## Results

### Chromosome X harbors little archaic coverage in humans today

We first review previously published estimates of archaic coverage on chromosome X and the autosomes in present-day human groups (Fig 1; 1–5). In all but one of the groups studied, chromosome X retains less archaic coverage than the autosomes. Only a Mandenka 1kG sample group has equal rates of archaic coverage on chromosome X and autosomes.

The autosomal to chromosome X coverage ratio reflects how many times greater the per-base pair rate of archaic coverage is on the autosomes than on chromosome X. The median coverage ratio considering all estimates is 7.91; excluding estimates specifically of Denisovan coverage [4], the median is 5.40.

Most estimates of archaic coverage displayed in Fig 1 are designed to identify coverage donated from Neanderthal specifically (ARGWeaver-D [3], Dical-admix [2], Sankararaman et al. [1], and Sankararaman et al. [4]), while Sankararaman et al. [4] also provides estimates specific to Denisovan ancestry (S2 Table). The method of Skov et al. [5] results in larger amounts of archaic coverage (compared to the other methods) likely because this method does not use a specific archaic donor to infer archaic ancestry. However, ratios of the autosome to chromosome X coverage rates computed from introgression maps inferred in Skov et al. [5] are comparable to the other methods.

Of the 120 human groups in which Denisovan coverage was identified, only 15 showed evidence of retaining any Denisovan coverage at all on chromosome X. Groups with Denisovan coverage on chromosome X were mostly found in Oceania and East Asia (S1 Table).

### Archaic coverage retention differs among continental groups

While chromosome X coverage rates are lower than autosomal rates in groups around the globe, we observe some broad patterns among continental groups.

The highest aut:chrX coverage ratios are found in some AMR 1kG sample groups, while some SAS 1kG sample groups (STU and BEB; Fig 2A, D) have the lowest autosomal to chromosome X coverage ratios. The high coverage ratios and relatively high variance in archaic coverage in these 1kG sample groups, particularly MXL and PUR, (see Table 1 for expanded names) may be due to recent admixture processes. (See S6 Fig and Discussion.)

**Fig 2.**
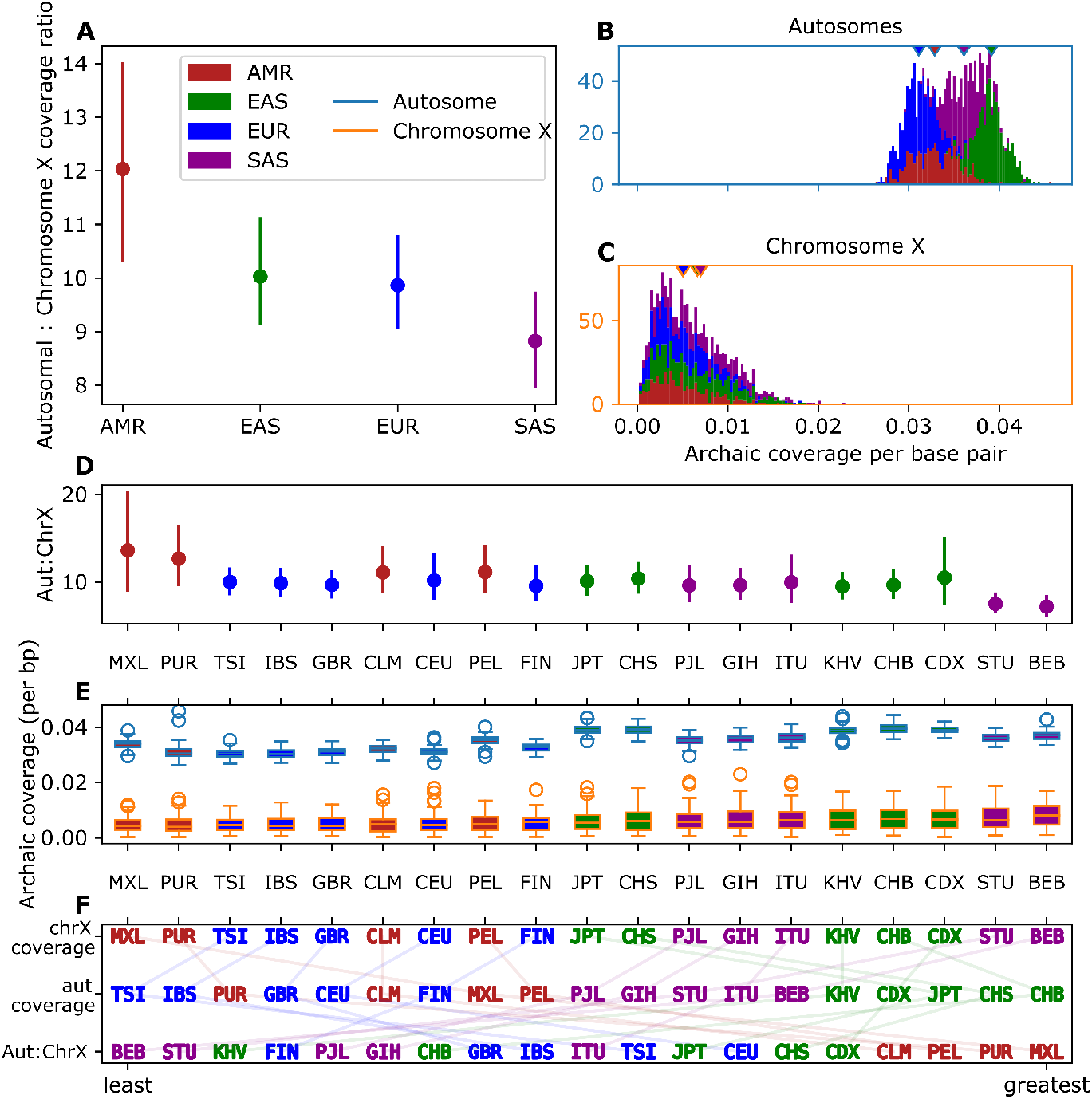
Distribution of archaic coverage between autosomes and chromosome X amongst 1kG samples. Coverage estimated by Skov et al. [5] from data from The 1000 Genomes Project Consortium [18]. See Table 1 for key to abbreviations. **A** Mean aut:chrX archaic coverage ratio among continental groups. Error bars show bootstrapped 95% confidence intervals around the means. **B** Distributions of archaic coverage on autosomes and **C** chromosome X across individuals, colored by 1kG continental group. Carets indicate means. **D** Summaries of coverage ratios, as well as **E** coverage distributions on the two chromosome types among 1kG sample groups. 1kG sample groups are sorted by ascending archaic coverage amounts on chromosome X. **F** Rank orderings of 1kG sample groups by mean chrX coverage, mean autosomal coverage, and aut:chrX ratio, sorted from least to greatest.

Differences among the coverage ratios are not cleanly correlated with absolute amounts of coverage. For instance, the EAS and EUR continental groups respectively retain the most and the least autosomal coverage (Fig 2B), yet exhibit middling and indistinguishable coverage ratio distributions (Fig 2A). Furthermore, more archaic ancestry on chromosome X does not imply more archaic ancestry on the autosomes. Ordering by each of the three metrics (mean chrX coverage, autosomal coverage, and aut:chrX ratio) produce different 1kG sample group-level rankings (Fig 2F).

### Dominance, sex-bias affect purging of archaic coverage from chromosome X

In order to study genomic patterns of archaic coverage, we implemented demographic simulations of a single pulse of introgression from a donor archaic population to the recipient modern human population (S1 Fig). Our simulation model closely follows that of Kim et al. [8]; complete simulation details are provided in Methods. For each simulation, we set the dominance model of mutations (all “additive” with *h* = 0.5, or all “recessive” with *h* = 0.0), and the sex-ratio of the introgressors (*p*, fraction of the introgressing individuals that are female). All mutations are deleterious, with most weakly so (≈ 74% of selection coefficients *s* ≤ 0.01). We generated autosomal and chromosomal haplotypes representing extant modern humans under the appropriate model of inheritance for each chromosome type. For each modern human haplotype sampled at the end of our simulation, we calculate its archaic coverage as the proportion of bases identical by descent to any of the introgressors, and contained within a contiguous tract of at least 500 bp.

Our simulations replicate the empirical observation of less archaic coverage on chromosome X than autosomes (Fig 3). Regardless of dominance model or sex-bias, the archaic coverage ratio between autosomes and chromosome X is greater than 1, indicating more autosomal than chromosome X archaic coverage.

**Fig 3.**
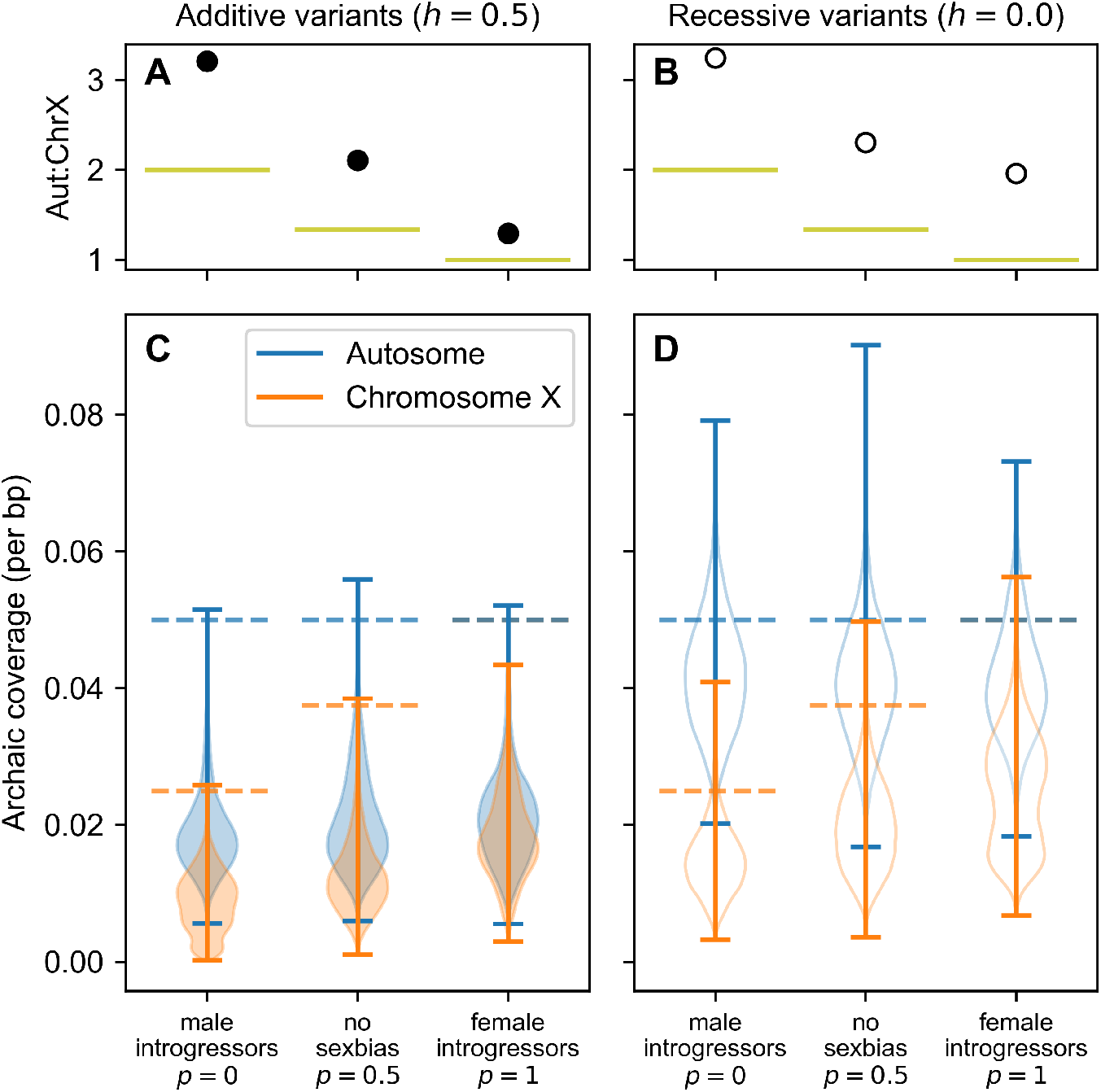
The effects of dominance and sex-bias on chromosome X archaic coverage. **A, B** Mean coverage ratios between a simulated autosome and chromosome X, estimated from 10,000 pairwise combinations of simulated autosomal and X chromosomes (see Methods). Bootstrapped 95% confidence intervals around these means are not visible because they are smaller than the markers. Filled points reflect additive variants (*h* = 0.5); hollow points reflect recessive variants (*h* = 0.0). Horizontal lines denote autosome to chromosome X ratio of archaic haplotypes in initial introgression pulse. See Eqn 1. **C, D** Filled violins depict additive variants; hollow violins depict recessive variants. Dashed horizontal lines denote magnitude of initial introgression pulse.

We also observe that archaic coverage is less than the initial introgression fraction of 5% in all models (Fig 3). Were archaic variants neutral in the modern human population, we would expect 5% archaic coverage to be maintained in the recipient population sample. However, the effective size of the archaic population in our simulations is smaller than that of the recipient modern human population. Deleterious variants that drifted to high frequency in the smaller archaic population are subsequently selected against after introgression into the larger modern human population, thereby ‘purging’ some of the initial introgressed sequences from the extant haplotypes. These results are consistent with previous studies of archaic purging by background selection [6, 7].

Under both dominance models, we observe a positive correlation between the female fraction of the introgressors and the rate of archaic coverage on chromosome X (Additive model: Pearson’s *ρ* = 0.53, *p <* 10^*−*20^; Recessive model: Pearson’s *ρ* = 0.37, *p <* 10^*−*20^; Fig 3). Autosomes are not inherited in a sex-specific fashion, so, as expected, we see a weaker relationship between archaic coverage and sex-bias on the autosomes (Additive model: Pearson’s *ρ* = 0.10, *p <* 10^*−*20^; Recessive model: Pearson’s *ρ* = *−*0.02, *p <* 10^*−*18^).

Introgression is an individual-level process. Therefore, the number of introgressing archaic haplotypes differs between autosomes and chromosome X depending on the sexes of the introgressors. We expect the ratio of autosomal haplotypes (*H*^*A*^) to X chromosomes (*H*^*X*^) to follow

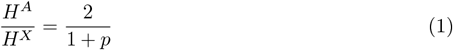

for an introgressing class with female fraction *p* (depicted as chartreuse lines in Fig 3A, B, and S2 Fig). For example, both the quantity and configuration of introgressing archaic haplotypes are identical between inheritance modes when all introgressors are female (*p* = 1; *i*.*e*. all chromosome X haplotypes introgress in pairs, just like autosomes). Indeed, we observe the smallest autosome to chromosome X coverage difference under the additive model when all introgressors are female (Autosomal mean 0.0211 vs. Chromosome X mean 0.0181; Fig 3C).

Thus, we both expect and observe a negative correlation between female fraction and the archaic coverage ratio between autosomes and chromosome X (Fig 3A, C, and S2 Fig). These findings are driven solely by the sex ratio of the introgressors and the inheritance model of the chromosome types, as our simulations do not invoke additional mechanisms such as chromosome-specific distributions of fitness effects, or hybrid incompatibilities.

However, not all of the difference between autosomal and chromosome X coverage can be attributed to the initial haplotype ratio within the introgression pulse. We observe autosome to chromosome X coverage ratios much higher than predicted by Eqn 1 in Fig 3A, C, particularly when *p <* 0.6 (S2 Fig). This effect is exacerbated when the sex ratio of the recipient population is male-biased; see for example S3 Fig).

The model of recessive variants produces greater coverage differences between autosomes and chromosome X than the model of additive variants across all degrees of sex-bias (Fig 3 and S2 Fig). This effect is driven by the heterotic advantage of archaic coverage in paired chromosomes when variants are recessive.

### Length distribution of archaic coverage tracts differs between chromosome X and autosome

Our simulations model recombination (using a variable recombination rate map derived from human data; see Methods Simulation procedure), and thus the lengths of the ancestry tracts can be informative of demographic events such as introgression [19]. We have access to the ground-truth ancestral history of each location on the simulated haplotypes, and can therefore identify whether any given base is identical by descent to any archaic introgressor, as well as identify the contiguous tracts of archaic coverage.

Although chromosome X harbors less archaic coverage than autosomes, its coverage is contained within longer tracts than autosomal coverage (Fig 4). This can be observed clearly in Fig 4C, which shows the proportion of all autosomal or all chromosome X tracts shorter than a given length, for all but the very longest tracts. Note that the lengths of the simulated chromosomes were the same and that the simulated autosome and chromosome X share the same recombination rate map and therefore the same mean recombination rate. (Although see S1 Appendix.) Differences between the autosome and chromosome X therefore reflect the different inheritance modes between the chromosome types.

**Fig 4.**
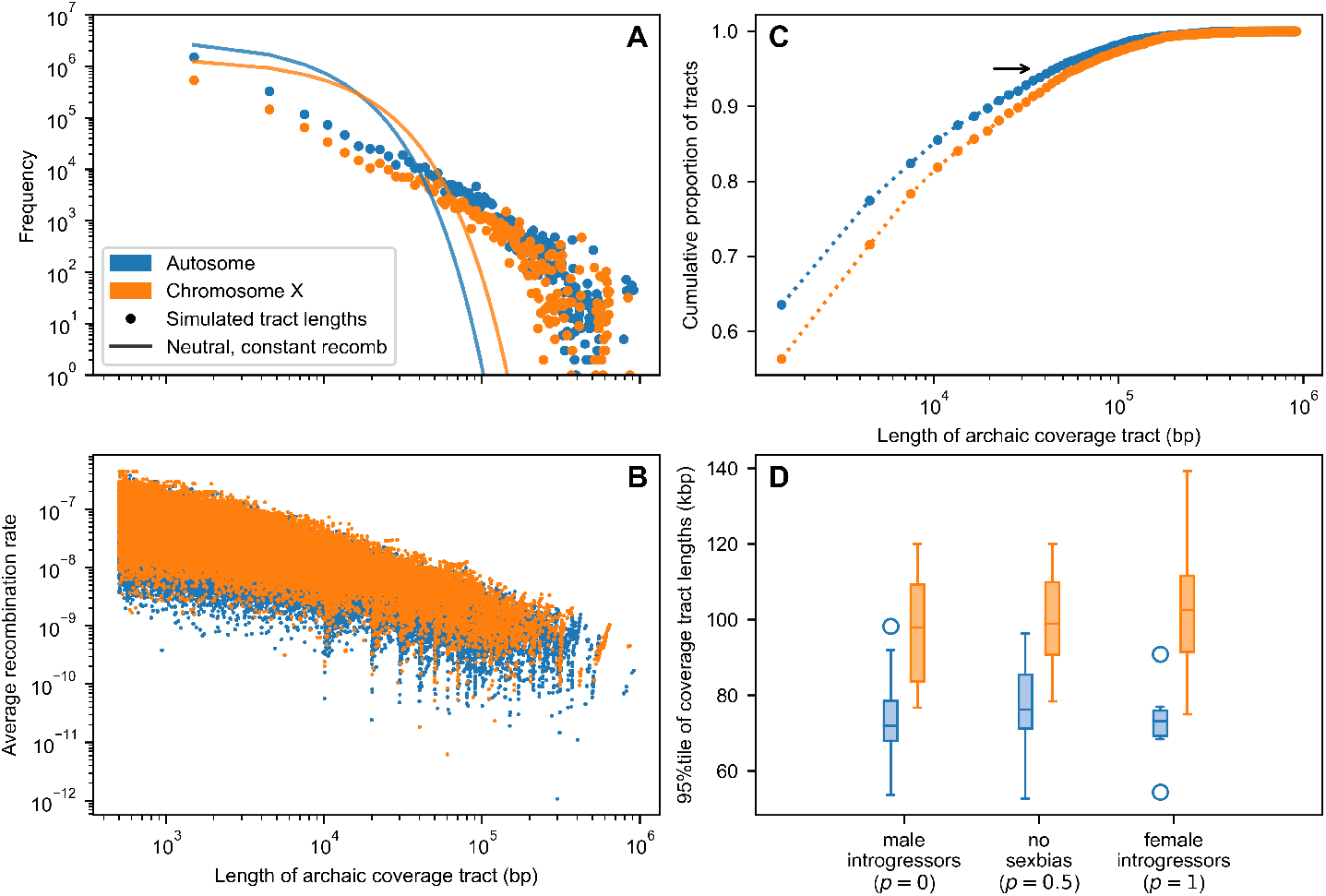
Archaic coverage tracts on chromosome X are longer than those on an autosome. All results shown reflect an additive model of dominance. **A** Tract length spectra from simulations of an archaic introgression scenario under autosomal or chromosome X inheritance; note log scale. Points indicate frequencies of archaic coverage tracts within length bins of 3000 bp. Simulated chromosomes shared a variable recombination rate landscape (see Methods for details). Solid lines indicate expected archaic coverage lengths under a simple, neutral model with constant recombination rate equal to the mean rate of the simulated recombination landscape. **B** Average recombination rate within each simulated archaic coverage tract, plotted against length of the tract. Note log scale. **C** Proportion of archaic coverage tracts that are a given length or shorter; proportions are calculated within each inheritance type. Arrow indicates 95th percentile of tract lengths, illustrated in panel D alongside sex-biased scenarios. **D** Distribution of 95th percentiles of archaic coverage tract lengths found on either an autosome or chromosome X, across degrees of introgressor sex-bias. Each box reflects the length distributions from ten simulation replicates.

As expected, due to the reduced frequency of recombination events in chromosome X inheritance and the preferential purging of archaic coverage from chromosome X, there are fewer total archaic coverage tracts on chromosome X than an autosome (Fig 4A). Compared to a simple model exponential model of coverage tract length decay from introgression to sampling (solid line in Fig 4A; see Methods), we find an excess of long tracts.

Panels A and B reflect data from simulations without introgressor sex-bias; panel C illustrates that regardless of introgressor sex-bias, the 95th percentile of archaic coverage tract lengths on chromosome X is longer than the 95th percentile of tract lengths on an autosome. Data shown is in the case of additive variants (*h* = 0.5), but this observation holds for the recessive case (*h* = 0).

Using a more stringent enrichment analysis, we find that the share of chromosome X coverage contained within the longest 1% of all tracts is greater than the share of chromosome X coverage within the remaining 99% of all tracts (S4 Fig). This pattern held for all additive models across all degrees of sex-bias. When variants were recessive and there were males among the introgressors (*p* = 0 and *p* = 0.5), this order was reversed.

### Coverage purging occurs on a fast timescale

In the preceding sections, we examined archaic coverage in extant haplotypes. We then studied the timecourse of archaic coverage retention from the introgression event to the sampling time.

Previous studies have observed that an equilibrium admixture quantity is achieved over a short timescale of tens of generations from the introgression event [7, 9, 20]. We recapitulate this result, and demonstrate its interaction with chromosomal inheritance type, dominance model, and sex-bias (Fig 5). We also observe a rapid temporal decay in the variance of archaic coverage across individuals (S5 Fig). In all models, mean introgression coverage falls halfway from its value in the first generation after introgression to its final, equilibrium value within 30 of the 15,000 intervening generations.

**Fig 5.**
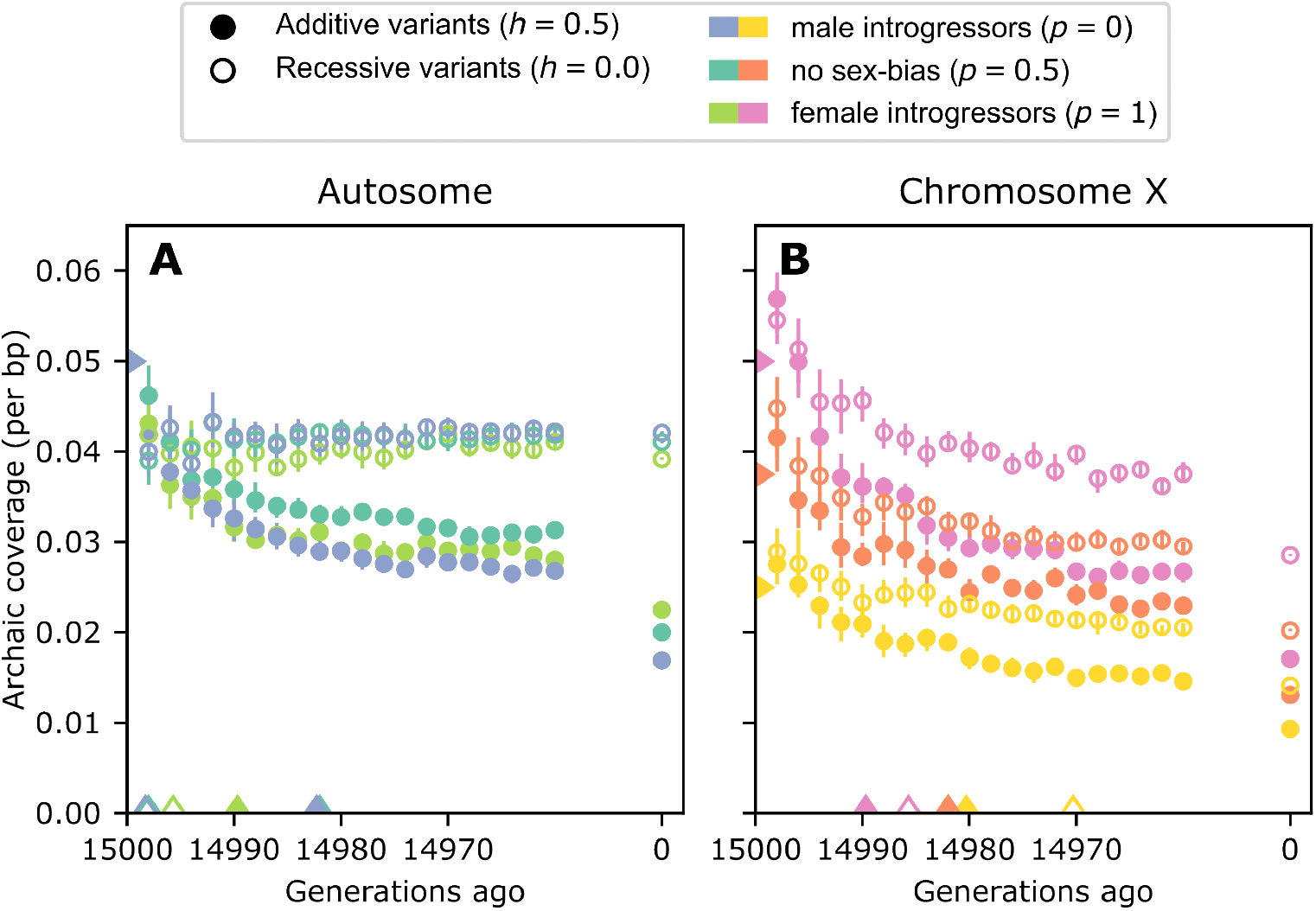
The rapid timecourse of archaic coverage purging on the autosomes and chromosome X. Carets on the x-axis indicate the generation at which mean coverage has fallen halfway to its final value from its value in the generation immediately following introgression. Carets on the y-axis indicate the initial introgression fraction. Error bars are bootstrapped empirical 95% confidence intervals around means of all samples across all simulation replicates. See S5 Fig for a visualization of the per-haplotype variance in archaic coverage over time. **A** Autosomal simulations. **B** Chromosome X simulations.

The timecourse of mean coverage in the autosomal recessive model is remarkably stable (Fig 5A).

As discussed, for the remaining scenarios (autosomal additive, chromosome X additive, chromosome X recessive), the final equilibrium coverage value is lower when more introgressors are male. This ordering by *p* emerges immediately on chromosome X, but is not observed until some time after the fortieth generation post-admixture in the autosomal additive model. Indeed, the greatest coverage differences between models are observed on chromosome X immediately after introgression (Fig 5B). Here, the dominance models are well-separated by sex-bias scenario. Within 20 generations, *p* no longer separates each curve, suggesting that sex-bias-related factors occur on a shorter timescale than those due to the dominance model of the variants.

## Discussion

Studies of archaic coverage in extant human genomes have repeatedly identified considerably less Neanderthal and/or Denisovan coverage on chromosome X than on the autosomes (Figs 1, 2, S1 Table). Several studies have attempted to explain this clear discrepancy by invoking hybrid incompatibility [1] or background selection [2, 6, 7]; the latter studies focused on background selection have posited that different distributions of selective effects on chromosome X and the autosomes explain observed data. Our study illustrates how sex-bias in demographic history, and specifically in the introgression event itself, can also generate lower archaic coverage on chromosome X than the autosomes. Although sex-bias has been suggested as an explanation for the deficiency of archaic coverage on chromosome X [2, 6, 8, 21], to our knowledge this is the first study to directly investigate how sex-bias can shape archaic coverage. We demonstrate that varying the sex-bias of the introgressors alone can significantly alter the aut:chrX ratio of archaic coverage. For example, under a model of additive deleterious variants, this ratio is more than doubled between a scenario with female introgressors and one with male introgressors (Fig 3A, S2 Fig).

### Heterosis

We also demonstrate the relative effects of models of dominance and sex-bias on the dynamics of archaic coverage purging from the autosomes and chromosome X (Fig 5). The effects of dominance and rate of purging have been examined on autosomes, without consideration of sex-bias, by multiple previous studies of archaic introgression [7–9]. Our simulation studies show that the heterosis is strongest when maintaining recessive archaic variants on the autosomes (Fig 5). This is expected on the autosomes, as heterosis maintains archaic coverage in admixed individuals [7, 22] by increasing heterozygosity at sites previously homozygous recessive for a deleterious allele. While heterosis helps maintain high archaic coverage on autosomes across time, on chromosome X the archaic coverage drops over time. This makes sense, as the hemizygous chromosome X in males can not benefit from increased heterozygosity.

### Aut:ChrX coverage ratios in 1kG samples

Examining the aut:chrX ratio of archaic coverage among 1kG continental groups reveals patterns distinct from continental group-level summaries of coverage amounts. As 2F demonstrates, all EAS and SAS 1kG sample groups contain more archaic coverage all than all AMR and EUR 1kG sample groups on both autosomes and chromosome X. However, ordering of aut:chrX ratio reveals a different pattern: all of the largest aut:chrX coverage ratios are found in AMR, and the rest of the ordering is a mix of EAS, SAS, and EUR 1kG sample groups.

Surprisingly, EAS and EUR have similar aut:chrX ratios, despite the absolute coverage amounts differing dramatically between them. Several studies have inferred distinct histories of archaic contact, including multiple introgression events [11, 23, 24]. If indeed the timing and number of archaic introgression events differ between EAS and EUR, it remains to be determined why their aut:chrX coverage ratios are so similar to each other.

The high coverage ratios and high variance in archaic coverage found in AMR 1kG sample groups, particularly the Mexican-American (MXL) and Puerto Rican (PUR) sample groups, may be due to recent admixture processes. (See S6 Fig for a visualization of ADMIXTURE [25] results of the AMR samples presented in Fig 2, using supervised clustering with *K* = 3 and EUR, AFR, and combined EAS and SAS samples as reference groups.) Recent admixture is evident in AMR 1kG sample groups, where African and Indigenous American ancestry is enriched on chromosome X in admixed individuals [26–28]. The AMR aut:chrX ratios therefore may display a higher spread due to the varying sources of their chromosome X coverage tracts.

### Future directions

In this work, our simulations follow the demographic model used in Kim et al. [8] (S1 Fig), which generates autosomal archaic coverage amounts on the order of these observed in humans today (S1 Table). We acknowledge that the true demographic histories are more complicated. Prior work on archaic introgression has mostly focused on quantifying archaic coverage, but we are only now starting to more fully characterize the introgression events: the timing of introgression, number of pulses, population size history, and how natural selection affects archaic variation in modern humans [11, 24].

Here, we simulate only deleterious variants, and these variants only accrue within exons. We also use the same variable recombination rate map for autosomal and chromosome X simulations (Figure 4; although see S1 Appendix). However, proximity to genes [1, 7], as well as differences in the local exon structure and recombination rate between chromosome X and autosomes [29, 30] may contribute to differences in the local distribution of archaic coverage [6, 8, 22, 31]. Future work may reveal the effect of drift and recombination on the quantity of archaic coverage that escapes background selection, and how these signals differ from positive selection around beneficial archaic variants [22].

Finally, we only consider parameters related to modern humans, but several studies have shown that introgression is common in other species, and characterizing its importance in speciation and adaptation are important questions that are currently being studied. Our model structure can be adapted to other organisms to provide insights into how demography, genomic structure, sex-bias and natural selection have shaped the distribution of introgressed variants on the autosomes and chromosome X.

Our work has demonstrated that dominance and sex bias affect the evolution of archaic coverage on chromosome X because of its unique inheritance pattern. We have shown that the observed low level of archaic coverage on chromosome X could be explained merely by a reduction in the effect of heterosis and sex-biases in the introgression events, without involving a more complex model with hybrid incompatibilities. Our work also suggests that negative selection was likely acting on archaic variants, and provides an appropriate set of null model for evaluating positive selection on introgressed segments on chromosome X. In general, by incorporating sex chromosomes explicitly into modeling frameworks, our work leverages more information from genetic data and begins to outline the set of evolutionary scenarios underlying the history and evolution of introgressed genetic variation.

## Materials and Methods

### Empirical coverage estimates

In Figs 1 and 2, we present published archaic coverage estimated from extant humans. References for the coverage estimates and data sources can be found in S2 Table. When necessary, we have converted all archaic coverage estimates to per-base pair rate of archaic genomic content, or the number of base pairs inferred to be of archaic origin, divided by either the total length chromosomes 1 through 22, or the length of chromosome X. Chromosome lengths were obtained from the GRCh37 assembly. Fig 2 presents archaic coverage estimated by the method of Skov et al. [5] in data from The 1000 Genomes Project Consortium [18]. We restrict the data to archaic coverage segments with posterior probability *>* 80%, in samples with calls on all of the autosomes and chromosome X. This way, the ratios of autosomal to chromosome X coverage are calculated within individual samples, not constructed by pairwise combinations of autosomal and chromosome X coverages (as in simulated data; see Simulation procedure). Data from 1835 individuals is presented; four individuals with autosomal to chromosome X coverage ratios greater than 200 have been removed. All code and data used to generate Figs 1 and 2 is either deposited with Dryad (DOI: 10.5061/dryad.xpnvx0kjq) or provided as Zenodo release XXX of the GitHub repository ramachandran-lab/archaic-chrX-sexbias.

### Simulation procedure

Forward genetic simulations under an explicit demographic model were implemented in the simulation software `SLiM v.3.3`[17]. Our simulation procedure and demographic model closely follow that of Kim et al. [8], although we have implemented sex-bias in the introgression pulse. For each simulation, we chose a model of chromosomal inheritance (autosomal or chromosome X), dominance (*h* = 0.5 or *h* = 0.0 for all mutations generated in the simulation), and introgressor sex-bias (fraction of introgressors that were female).

### Genomic model

Our genomic model is identical to that described in the *Simulations of human genomic structure* section of Kim et al. [8], with the exception that our simulations used a scaling factor of 2, rather than 5.

Inheritance of chromosomes via either an autosomal or chromosome X pattern was managed by SLiM. Each simulated chromosome was 100Mb in length, with identical exon placement and recombination rate maps. The choice of exonic regions and variable recombination rate maps is described in Kim et al. [8]. See S1 Appendix for a sketch of Fig 4 results with different exon placement and recombination rate maps between chromosome X and autosomal simulations.

Chromosomes accumulated mutations within exonic regions. Mutations had selection coefficients drawn from a distribution fit to data from European individuals (EUR population, The 1000 Genomes Project Consortium [18] Phase 3; see Kim et al. [32]) of the form:

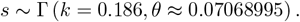

### Demographic model

We simulated a simple two-population model with a single-generation pulse of introgression from source to recipient (S1 Fig).

Simulations reflected a 25,000-individual recipient population, which received a pulse of introgression 5% of this size from a 250-individual source population after 20,000 generations of isolation. Simulation was terminated 15,000 generations after the introgression pulse. Simulations were scaled down by a factor of 2 for computational efficiency, following best practices [17]. Following simulation in SLiM, a joint, pre-isolation history was appended to the simulation using the `recapitate()`method provided by `pyslim v.0.403`. All code necessary to perform the simulations is provided as Zenodo release XXX of the GitHub repository ramachandran-lab/archaic-chrX-sexbias.

### Introgression coverage calculations

Once a simulation was complete, we calculated introgression coverage in each of 1000 haplotypes sampled from the final generation of individuals in the recipient population. A haplotype’s coverage (*c*_*h*_) is the proportion of its bases that are identical by descent (IBD) to one or more of the introgressing archaic haplotypes (*H*^*A*^). Coverage is calculated as:

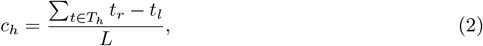

where *L* is the length of the haplotype in base pairs, and

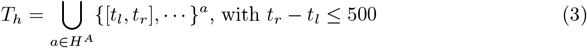

where *t*_*l*_, *t*_*r*_ are the left and right endpoints of a tract of haplotype *h* that is shared IBD with introgressing archaic haplotype *a*.

Haplotype sampling and IBD analysis for each simulation replicate were performed on the tree sequence data structure output from SLiM. The locations of the tracts shared IBD were obtained from the tree sequence using the `find_ibd()`method implemented in `tskit v.0.3.2`[33].

Due to limitations of the simulation software, simulations of autosomal inheritance and chromosome X inheritance were performed separately. Therefore, there is no natural pairing of particular autosomal and chromosome X haplotypes under which to calculate an aut:chrX coverage ratio. The simulated aut:chrX ratios presented in Figs 3AB, S2 Fig, and S3 Fig is the mean and 95% confidence interval of 2000 bootstrap resamplings of the ratio of *n* choices of autosomal coverages and *n* choices of chromosome X coverage, chosen with replacement from all coverage data (which each contained at least *n* unique haplotypes). For Fig 3, *n* = 10, 000.

All code and data used to calculate coverage amounts and tract lengths, as well as plot Figs 3, 4, and 5, is either deposited with Dryad (DOI: 10.5061/dryad.xpnvx0kjq) or provided as Zenodo release XXX of the GitHub repository ramachandran-lab/archaic-chrX-sexbias.

## Supporting information

**S1 Appendix. Description of additional simulations with chromosome-specific exon and recombination maps**.

## Acknowledgments

We gratefully acknowledge productive discussions with members of the Ramachandran and Huerta-Sánchez labs, particularly David Peede.

## Supplementary Material

## S1 Appendix

We performed additional simulations with chromosome-specific exon and recombination maps. Simulations of realistic chromosome architecture used different exon and recombination rate maps for simulated autosomal and chromosome X chromosomes.

These simulations modeled a randomly-selected 50Mb portion of the long arm of either chromosome 7 (`chr7:101351049-151351049`) or chromosome X (`chrX:101351049-151351049`). Bulk properties of these regions are presented in Table S3.

Exon locations were defined as the protein coding regions in the basic gene annotation map obtained from GENCODE Release 37 (GRCh37; https://www.gencodegenes.org/human/release_37lift37.html) [38].

Recombination rate maps were obtained from Bhérer et al. [30], and reflect data from individuals with European ancestry. We used a sex-averaged map for chromosome 7 simulations.

---

**Fig S1.**
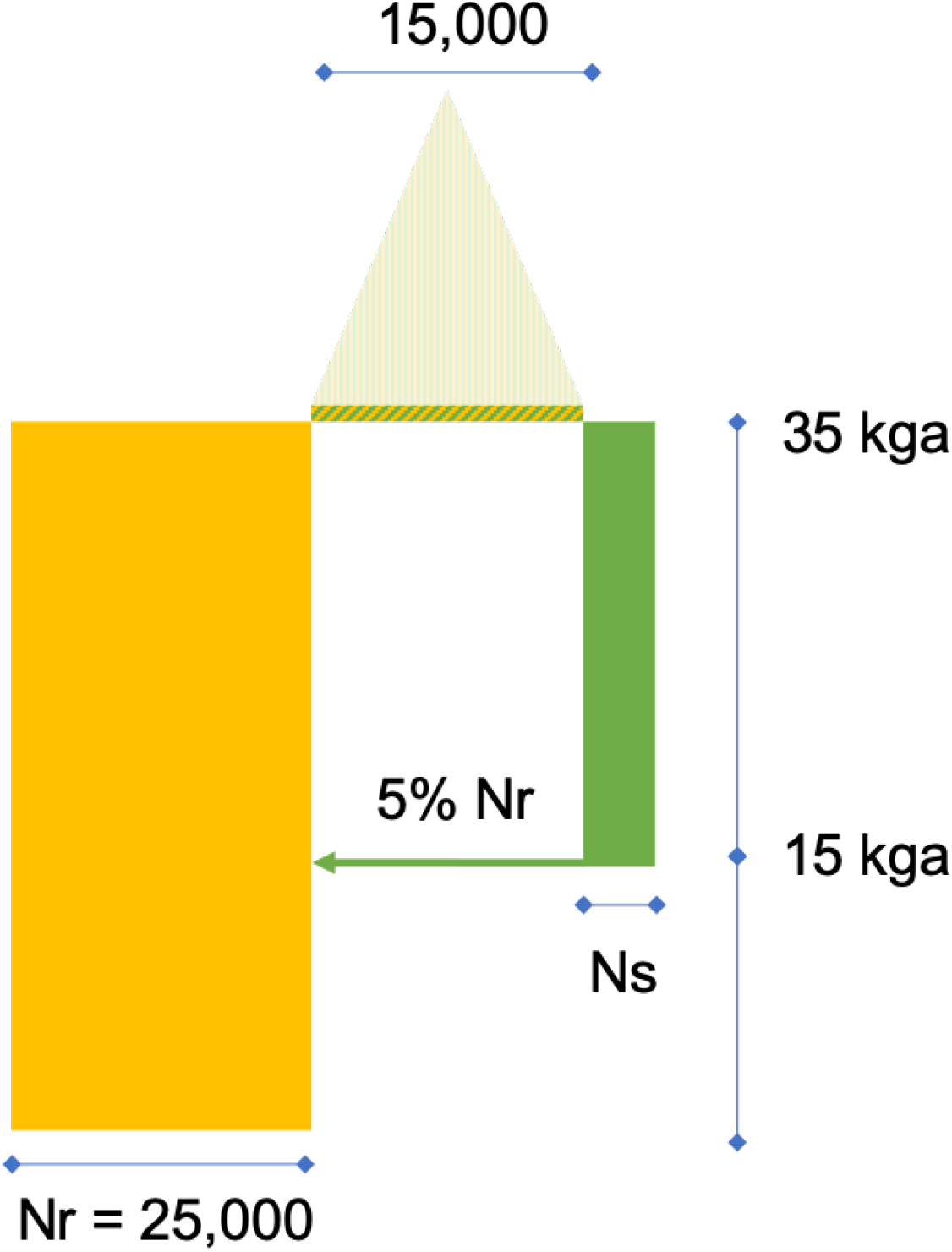
Cartoon of demographic model implemented in simulations. *N*_*r*_ indicates size of recipient population in haplotypes; *N*_*s*_ indicates size of source population in haplotypes; *kga* indicates thousand generations ago. Results presented here have *N*_*r*_ = 100*N*_*s*_. Filled colors indicate generations in forward simulation; a prior coalescent history is appended to the ancestral population. See Methods for further details.

**Fig S2.**
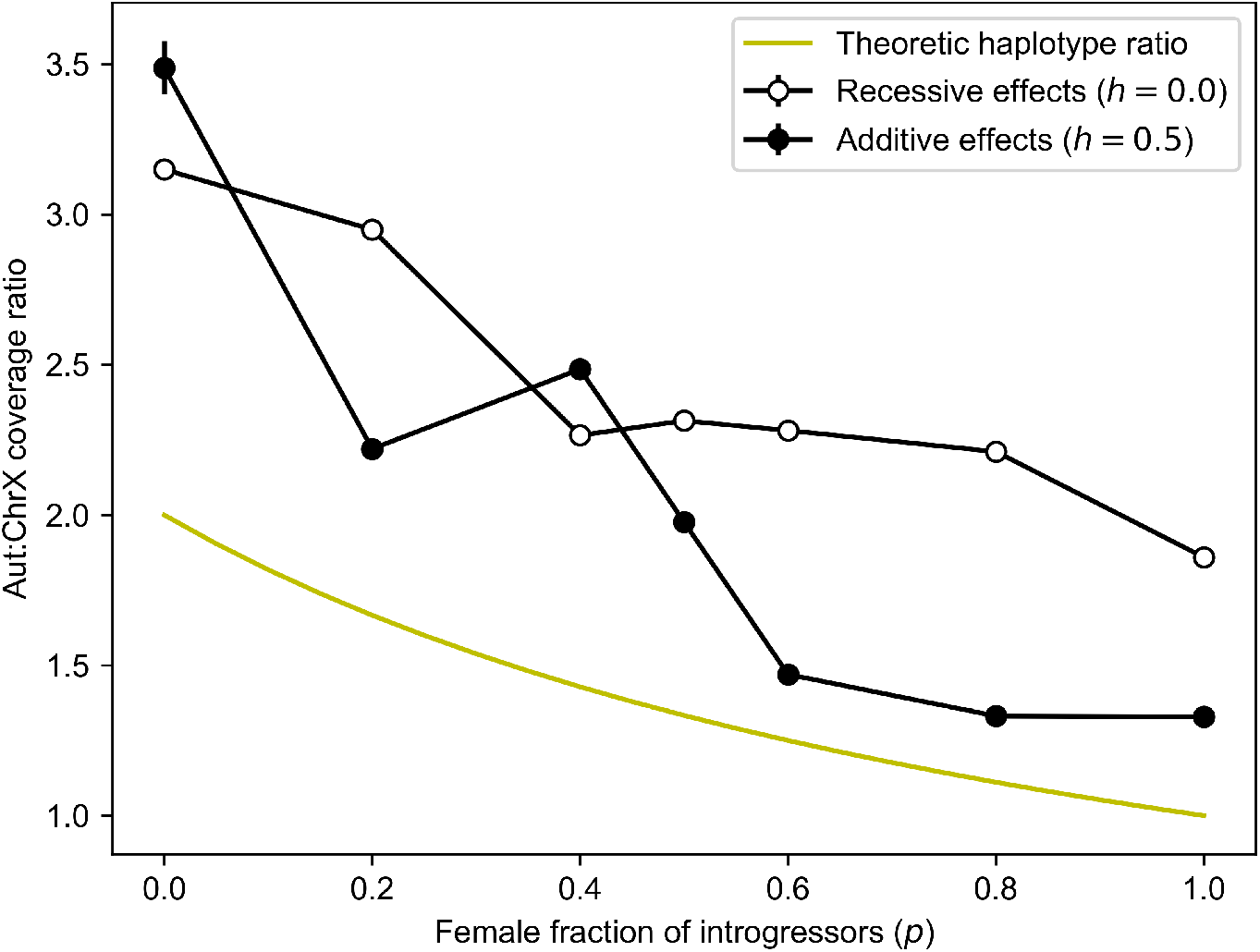
Autosome to chromosome X coverage ratio decreases with greater female fraction of introgressors in simulations. Chartreuse line depicts autosome to chromosome X haplotype ratio of a theoretical population with a given male fraction (Eqn 1). Error bars are bootstrapped 95% confidence intervals around the mean of 10,000 ratios generated from autosomal and chromosome X coverage distributions.

**Fig S3.**
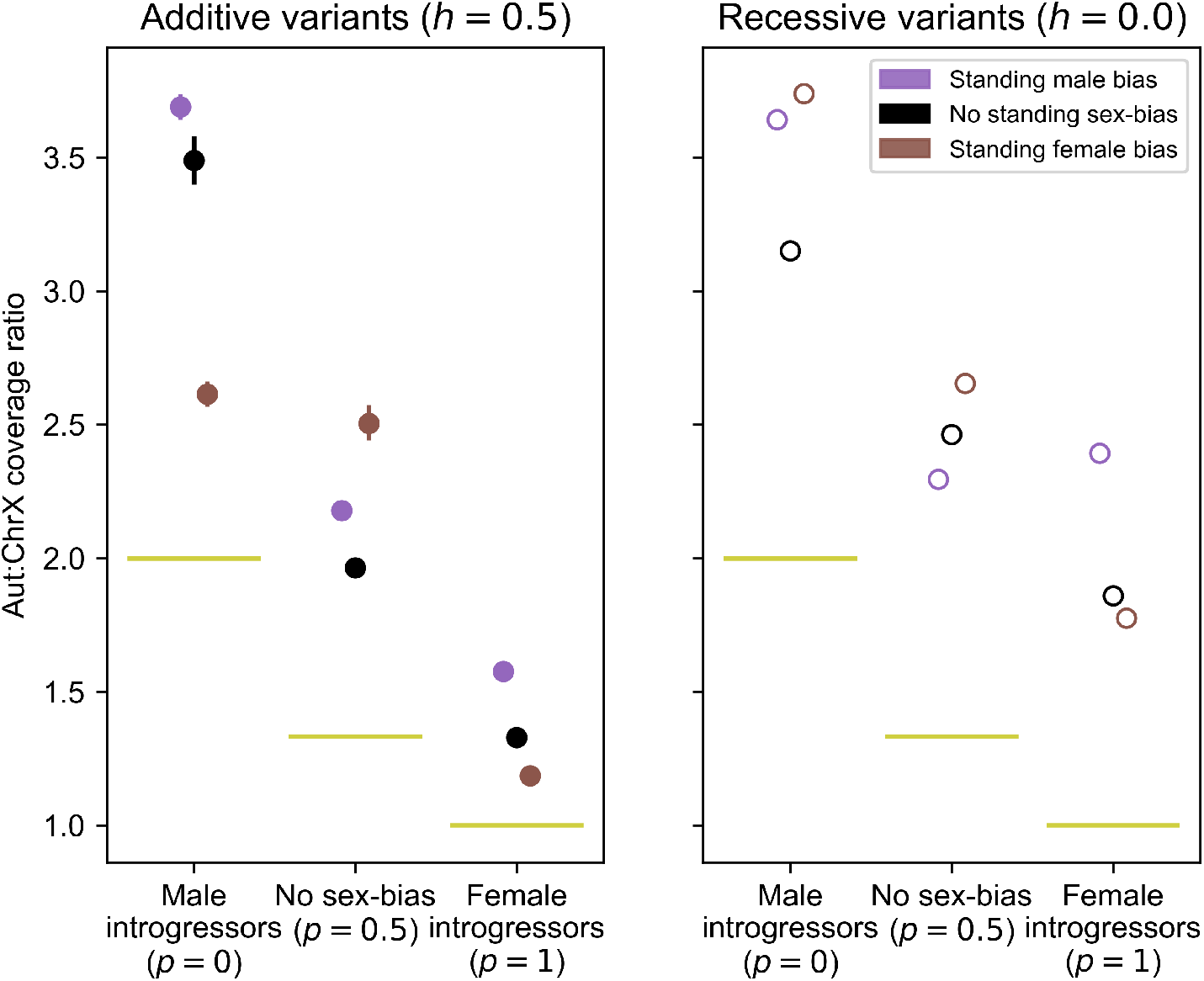
An unequal sex ratio in the recipient population interacts with introgressor sex-bias. **A** Aut:ChrX coverage ratios from simulations that reflect both sex-bias within the introgression pulse, and a constant, (“standing”) unequal sex ratio in the recipient population. Sex ratios shown are 25% female (standing male bias), 50% female (no standing sex-bias), and 75% female (standing female bias). Black points reflecting no standing sex-bias are equivalent to those shown in Fig 3A and C. Error bars are bootstrapped 95% confidence intervals around the mean of 10,000 ratios generated from autosomal and chromosome X coverage distributions; in most cases the error bars are smaller than the displayed points. All variants are additive (*h* = 0.5). **B** Same as panel A, but all variants are recessive (*h* = 0).

**Fig S4.**
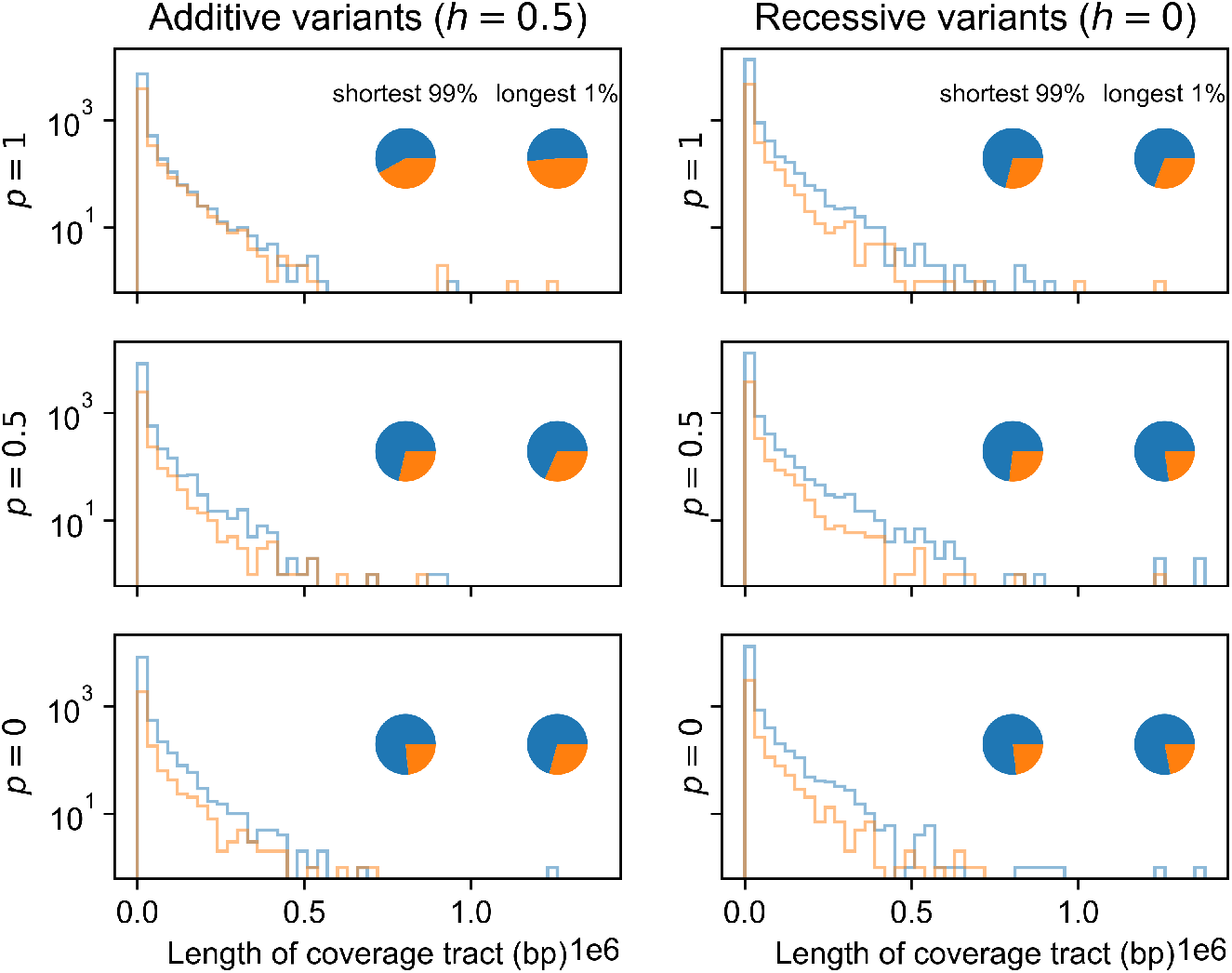
The longest archaic coverage tracts are preferentially found on chromosome X in our simulation studies. Archaic coverage tract length spectra from a simulated autosome (blue) and chromosome X (orange). Each plot represents a combination of dominance (*h* = 0 or *h* = 0.5) and sex-bias (*p* = 0, *p* = 0.5, or *p* = 1, where *p* is the female fraction of the introgressors). Pie charts depict fraction of total coverage found on the autosome or chromosome X. Right pie shows coverage contained within the longest 1% of tracts; left pie shows all other coverage. Note that chromosomes have identical size and local recombination rates; see Methods.

**Fig S5.**
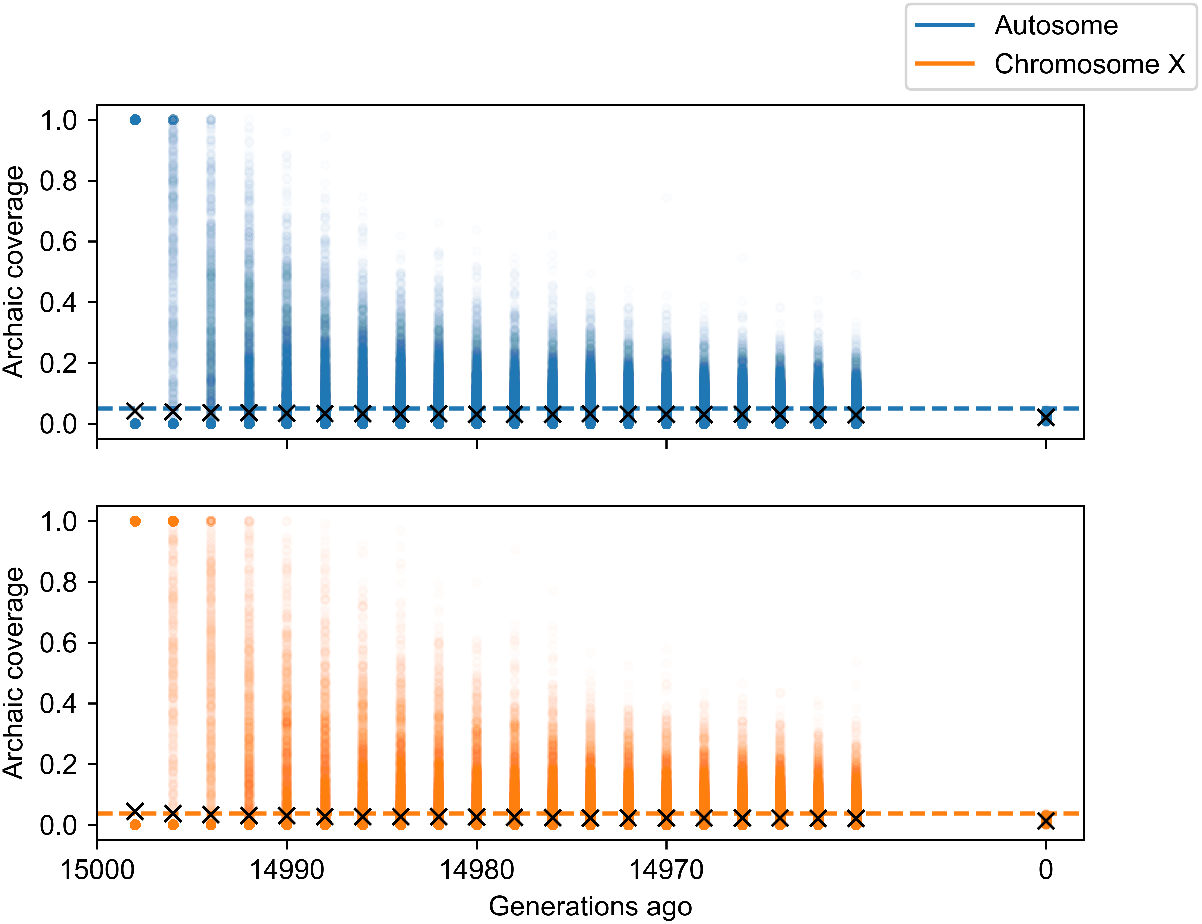
Variance in archaic coverage per haplotype decays rapidly after introgression. Each point (note transparency) reflects the archaic coverage on one of 1000 haplotypes sampled from the recipient population every two generations for the first 40 generations after the introgression event. Crosses indicate mean coverage at each timepoint (see Fig 5). Horizontal line indicates initial introgression fraction.

**Fig S6.**
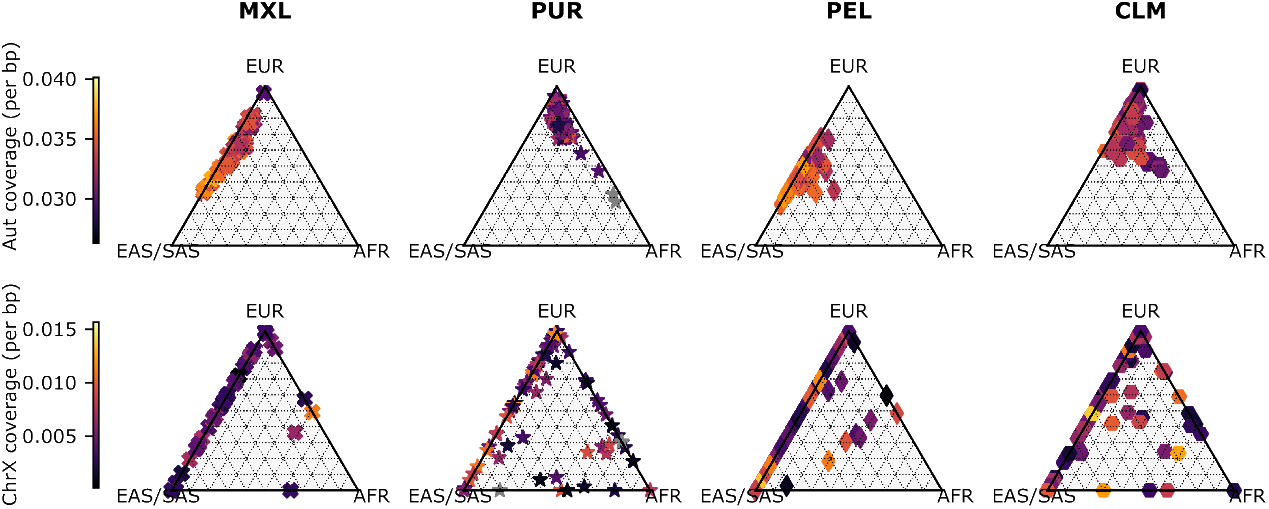
Autosomal and chromosome X archaic coverage varies among AMR 1kG sample groups. Colors indicate per-base pair archaic coverage estimates on autosomes or chromosome X, inferred by method of Skov et al. [5] using data from AMR samples from The 1000 Genomes Project Consortium [18], selected as described in Methods Empirical coverage estimates. Coordinates are ADMIXTURE [25] ancestry estimates for the individuals, using supervised clustering with K=3 and EUR, AFR, and combined EAS and SAS individuals as reference groups. PUR samples HG01108 and HG01241 have exceptionally high autosomal archaic coverage (0.046 and 0.042, respectively), and so are marked in grey to preserve color scaling.

**Fig S7.**
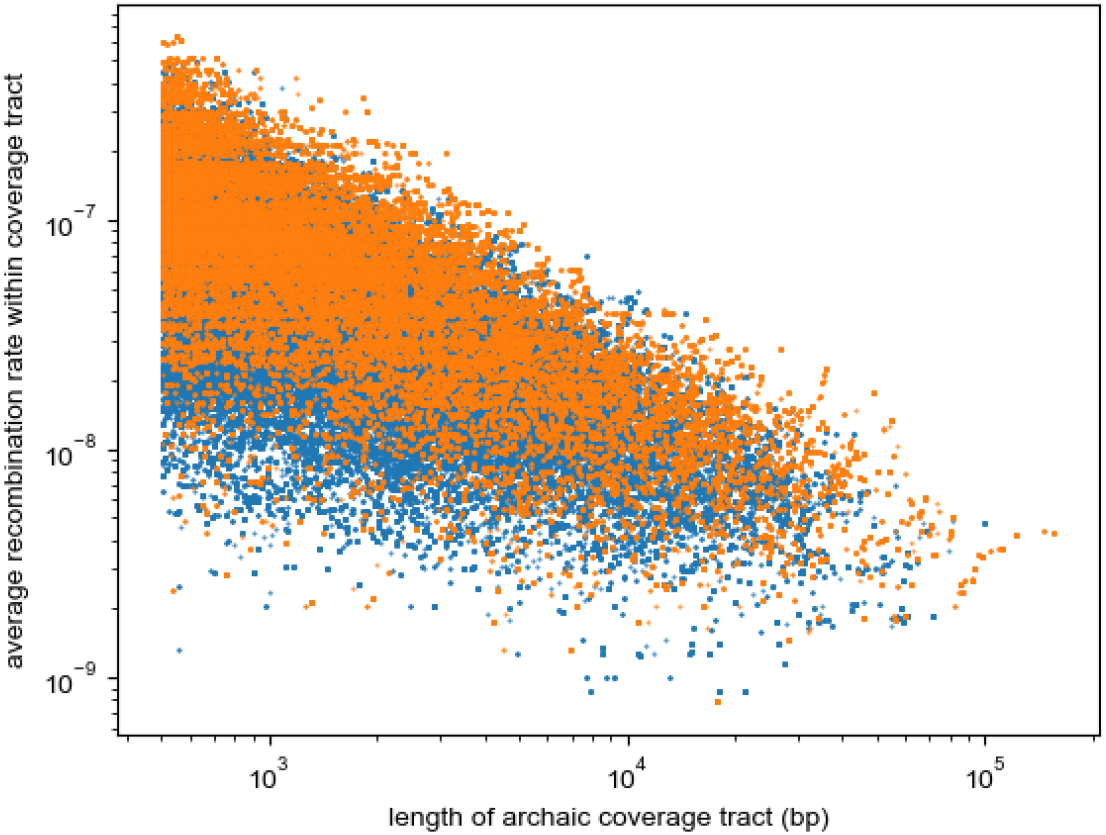
Recombination rate and length of archaic coverage tract are negatively correlated. Identical to Fig 4B, plotted with data from chromosome-specific exon and recombination rate maps as described in S1 Appendix.

**Table S1.**
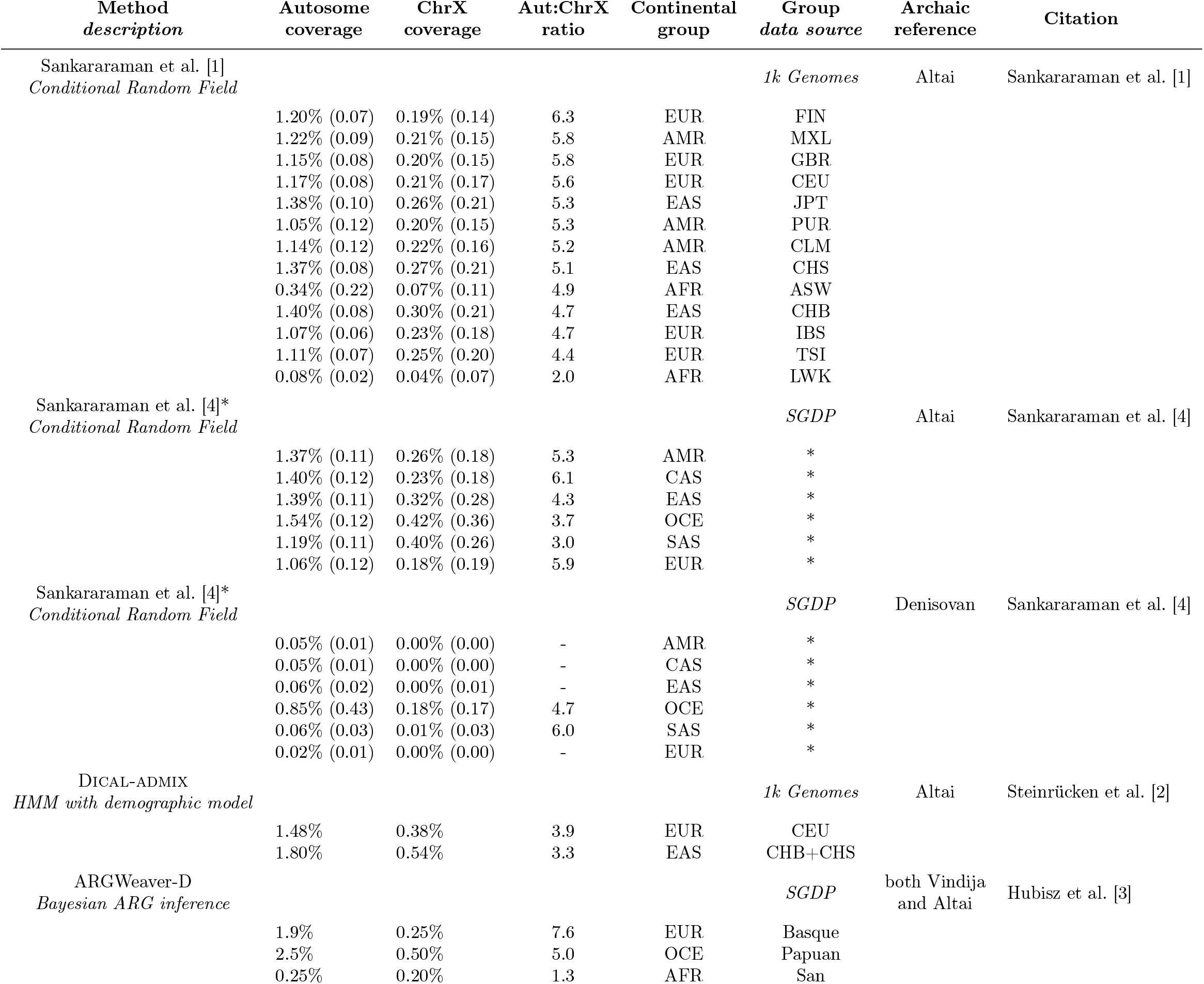
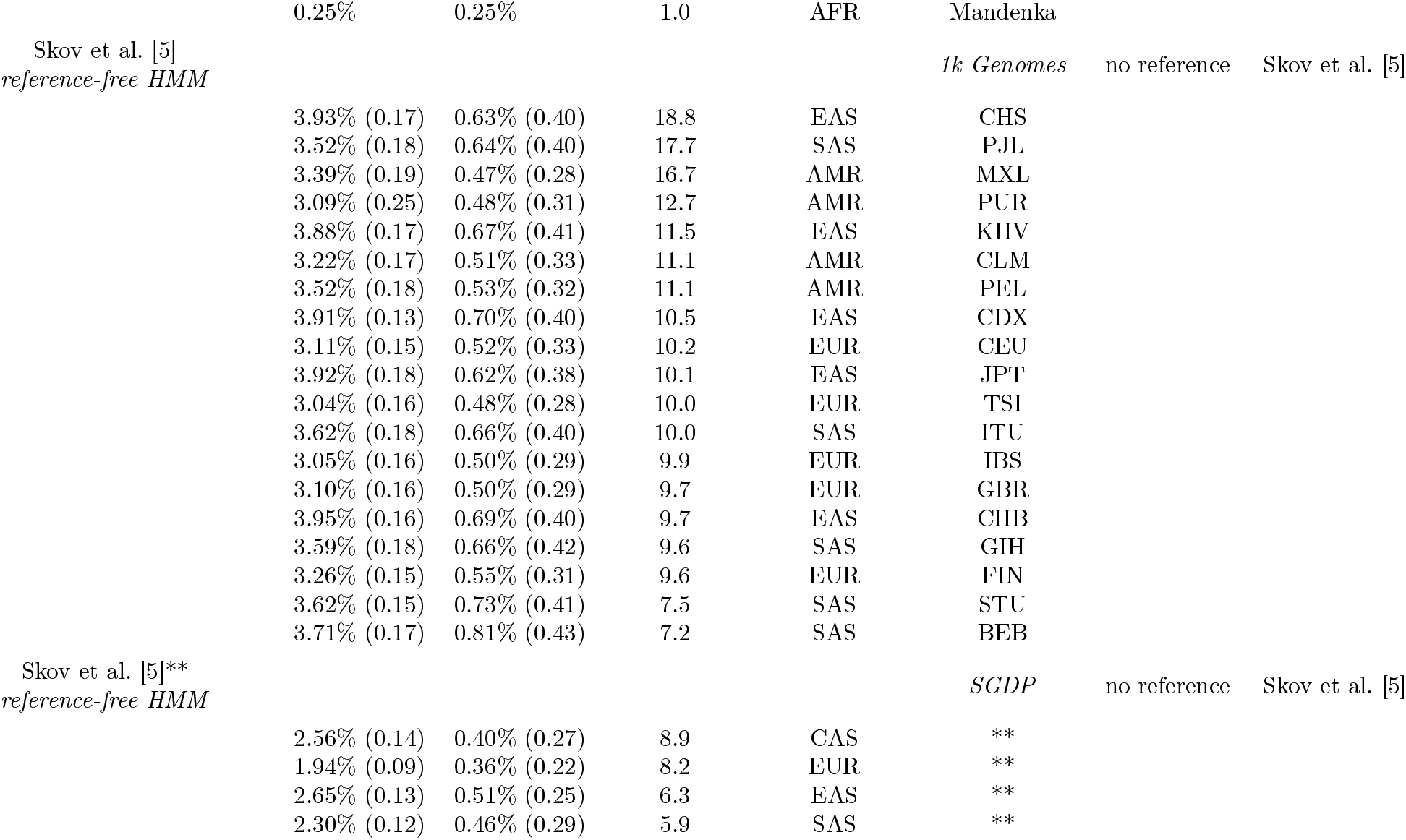
Genome-wide archaic coverage levels on autosomes and chromosome X (chrX) inferred from human genomic data. Coverage is the per-base pair proportion of the genome inferred to have come from archaic sources, found either on the combined 22 autosomes, or on chromosome X. The standard deviation across individuals is in parentheses. The fourth column shows the ratio of autosomal archaic coverage to chromosome X archaic coverage (Aut:ChrX), which is always at least equal, and can range up to 18.8. All groups outside of Continental Africa have at least 3.3 times more relative coverage on autosomes than chromosome X. See Table 1 for continental group and 1kG sample group abbreviations. Altai Neanderthal published in Prüfer et al. [34]; Vindija Neanderthal published in Prüfer et al. [35]; Denisovan published in Meyer et al. [36]. Inference methods were applied to various data sources: The 1000 Genomes Project Consortium [1k Genomes; 18] and Mallick et al. [SGDP; 37]. * See Supplementary Table 2 of Sankararaman et al. [4] for group-specific coverage estimates. ** See Supplementary Dataset S5 of Skov et al. [5] for group-specific coverage estimates.

**Table S2.** Methods used to estimate archaic coverage on chromosome X. Coverage estimates in extant human groups are presented in Fig 1.

| Fig 1 label | Estimation method | Data Source* | Archaic specificity |
| --- | --- | --- | --- |
| ARGWeaver-D | Hubisz et al. [3] | SGDP | Neanderthal |
| Dical-admix | Steinrücken et al. [2] | 1kG | Neanderthal |
| Sank14 | Sankararaman et al. [1] | 1kG | Neanderthal |
| Sank16Neand | Sankararaman et al. [4] | SGDP | Neanderthal |
| Sank16Denis | Sankararaman et al. [4] | SGDP | Denisovan |
| Skov1kG | Skov et al. [5] | 1kG | none |
| SkovSGDP | Skov et al. [5] | SGDP | none |

\*SGDP refers to Simons Genome Diversity Project [37]; 1kG refers to The 1000 Genomes Project Consortium [18].

**Table S3.** Properties of simulated regions of realistic human chromosome architecture.

| Inheritance model | Avg. Exon Density | Avg. Recombination Rate (cM/bp) |
| --- | --- | --- |
| Autosomal | 0.026 | $1.3 \times 10^{-7}$ |
| Chromosome X | 0.019 | $2.2 \times 10^{-7}$ |

